# Unraveling the Impact of Gene Length on Kinetic Parameters: Implications in Drug Target selection

**DOI:** 10.1101/2024.08.31.610572

**Authors:** Soham Choudhuri, Bhaswar Ghosh

## Abstract

Gene expression is a multifaceted process crucial to understanding molecular biology and pharmacology. Our research focuses on elucidating the intricate relationship between gene length and kinetic parameters, such as *S*_*i*_, *K*_*on*_, *K*_*off*_, and *SK*_*off*_, which significantly influence the mean expression levels of genes.Using a two-state stochastic gene expression model implemented in Python, we analyzed single-cell transcriptomics data to predict kinetic parameters for each gene. We classified genes into short and long categories, revealing distinct patterns in the relationship between gene length and these parameters. Our results indicate that burst size plays a critical role in mean expression, highlighting its importance for identifying gene targets that require lower drug doses for therapeutic effects.

## Introduction

Understanding the intricate dynamics of gene expression is a cornerstone of molecular biology and pharmacology. Gene expression is a complex process influenced by a multitude of factors, from transcriptional initiation to post-translational modifications ***Gibney and Nolan (2010)***. Each step in this cascade is finely tuned, shaping the cellular landscape and ultimately influencing physiological outcomes ***Li et al. (2022)***. Our research has explored the relationship between gene length and various kinetic parameters, identifying which parameters are most crucial for the mean expression level of a gene. Additionally, we have investigated strategies for selecting genes that require lower drug doses for therapeutic effects.

To support this research, we have written Python script for a two-state stochastic gene expression model ***Kim and Marioni (2013)***. By inputting single-cell transcriptomics data, this model predicts the values of different kinetic parameters for each gene ***Kim and Marioni (2013)***. Next, we find the relationship between gene length and different kinetic parameters. We have divided gene lengths into two categories, long and short. We have shown the relationship pattern between gene length and different kinetic parameters.

Understanding the impact of various kinetic parameters on mean expression is essential for uncovering the mechanisms of gene regulation and transcription dynamics ***Volteras et al. (2023)***. Kinetic parameters such as transcription rate (*S*_*i*_), *K*_*on*_, *K*_*off*_, Burst size (*SK*_*off*_) play critical roles in determining how genes are expressed under different conditions ***Choi et al. (2017); Yu et al. (2023)***. *S*_*i*_ is Transcription rate, Determines the speed at which RNA polymerase synthesizes RNA from DNA. A higher rate means more RNA transcripts and potentially more protein production ***Lodish (2008)***. *K*_*on*_ is Association rate, Reflects how quickly transcription factors bind to DNA to initiate transcription. A higher *K*_*on*_ can lead to more frequent transcription initiation. *K*_*off*_ is Dissociation rate, indicates how quickly transcription factors detach from DNA. A lower *K*_*off*_ means transcription factors remain bound longer, potentially leading to more transcription. *SK*_*off*_ is burst size, represents the number of RNA transcripts produced in a single burst when transcription factors are bound ***Baron and van Oudenaarden (2019)***. Larger bursts can result in higher levels of gene expression. By studying these kinetic parameters, researchers can gain insights into the regulatory elements and factors that control gene expression. We have used linear regression model and showed that burst size has more impact on the mean value of gene expression.

Burst size and mean value expression of gene are two important factor that can help to choose target with lower dose. Choosing drug targets that necessitate lower doses of drugs can reduce the potential for side effects, thereby enhancing patient compliance and quality of life ***Sorger et al. (2011)***. Lower doses also improve safety by minimizing toxicity concerns, which is especially important for long-term therapies or vulnerable populations ***Kulkarni (2016)***. Economically, lower dosages translate to reduced production and shipping costs, making treatments more accessible and affordable. Additionally, simplified dosing regimens with fewer adverse effects can lead to better treatment outcomes and increased patient adherence. In the context of antibiotics and cancer therapies, using lower doses can also help prevent the development of resistance, contributing to more sustainable and effective healthcare practices ***Spellberg and Gilbert (2014)***.

## Results

Gene expression is a complex phenomenon influenced by a wide range of factors. From transcriptional initiation to post-translational modifications, each step in this cascade is finely tuned, shaping the cellular landscape and ultimately influencing physiological outcomes ***Lodish (2008)***. Our work has shed light on the relationship between gene length and kinetic parameters, impact of kinetic parameters on mean value expression and selecting drug target for which lower dose is needed. We have used a stochastic model of gene expression ***Kim and Marioni (2013)***. The steadystate solution of this model is a hypergeometric function. We can’t directly compute this solution. So, some assumptions has been made to simplify the steady-state result. Finally, we get a poisson-beta distribution. A hierarchical Bayesian model has been used to estimate the parameters of the Poisson-beta distribution. Using Gibbs sampling, we get different kinetic parameter values from single-cell transcriptomics data used as input in this model ***Kim and Marioni (2013)***. We have implement this mathematical model in Python. The main advantages of using Python was the simplicity of the language, and the plethora of libraries available. A language like C++ could also be used, but it would require us to implement things like correlation and covariance ourselves. Hence we found Python to be more suitable. We classified genes into short and long categories, revealing distinct patterns in the relationship between gene length and these parameters. Our results indicate that burst size plays a critical role in mean expression, highlighting its importance for identifying gene targets that require lower drug doses for therapeutic effects. We have given a detail discussion on this results.

### Relationship pattern between length and other kinetic parameters

We have found the relationship between the length of genes and kinetic parameters. We have divided the gene lengths into long and short lengths. We have seen the Long and short length genes follow different pattern with same kinetic parameter. Relationship pattern between short length and kinetic parameters can be shown in figure 2. Relationship pattern between long length and kinetic parameters can be shown in figure 3. We can see from the figure 2, 3 that *K*_*on*_ and *K*_*off*_ make cluster in a region for all different length. We are getting decreasing pattern of S_*i*_ as we increase the genelength. Longer genes generally take more time to be transcribed compared to shorter ones. This is because RNA polymerase must travel along the entire length of the gene to synthesize the mRNA, which naturally takes longer for longer sequences ***Roeder (2005)***. The efficiency of transcription can be influenced by gene length. Longer genes may be transcribed less efficiently due to the increased likelihood of encountering obstacles, such as DNA-bound proteins or chromatin structures, which can impede the progress of RNA polymerase ***Kong and Lasko (2012)***. The architecture of a gene, including the presence of introns and exons, can also affect transcription. Longer genes often have more introns, which need to be spliced out, potentially affecting the overall transcription process and subsequent mRNA processing ***Tian and Manley (2017)***. So, longer genes generally require more time to be transcribed, various factors, including transcriptional regulation, gene architecture, and the cellular context, can modulate how gene length affects the rate of transcription.

**Figure 1.**
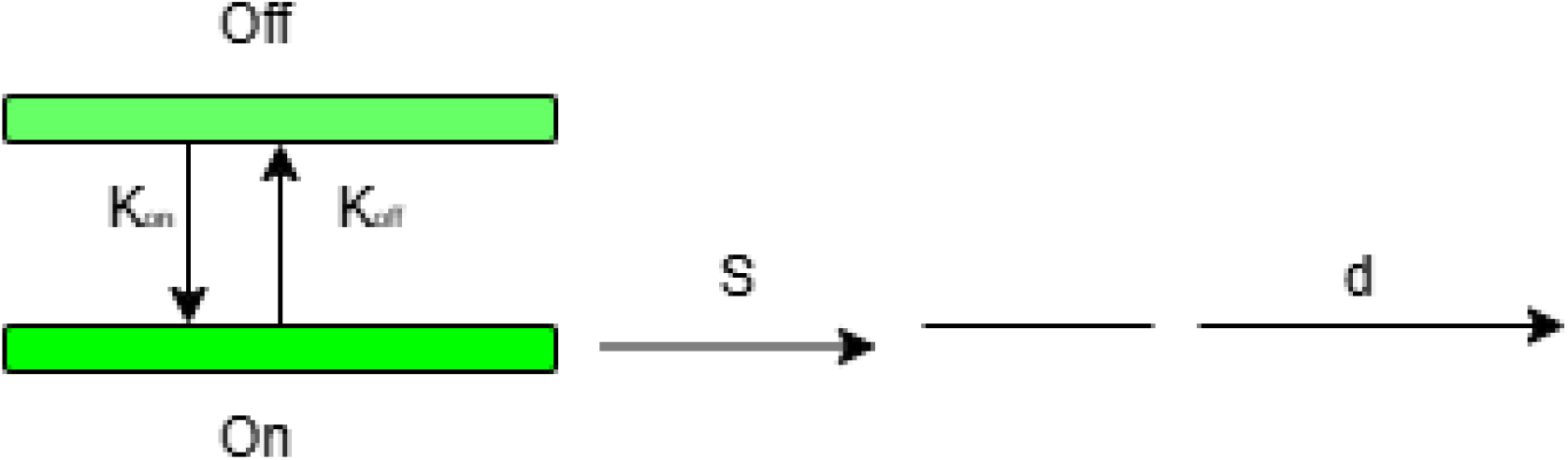
Schematic of a two-state kinetic model for stochastic gene expression

**Figure 2.**
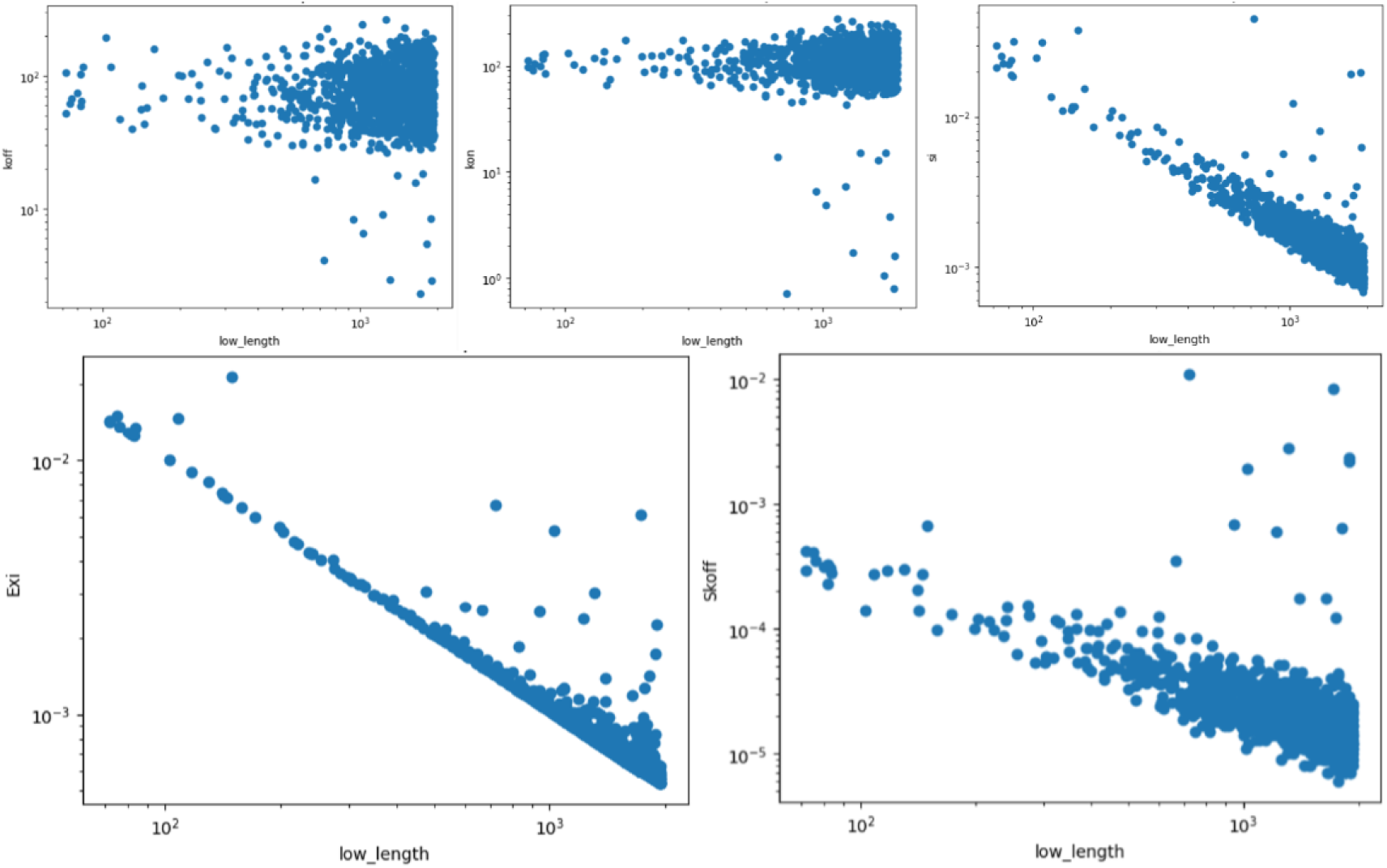
Relationship pattern between Short length genes and kinetic parameters.

**Figure 3.**
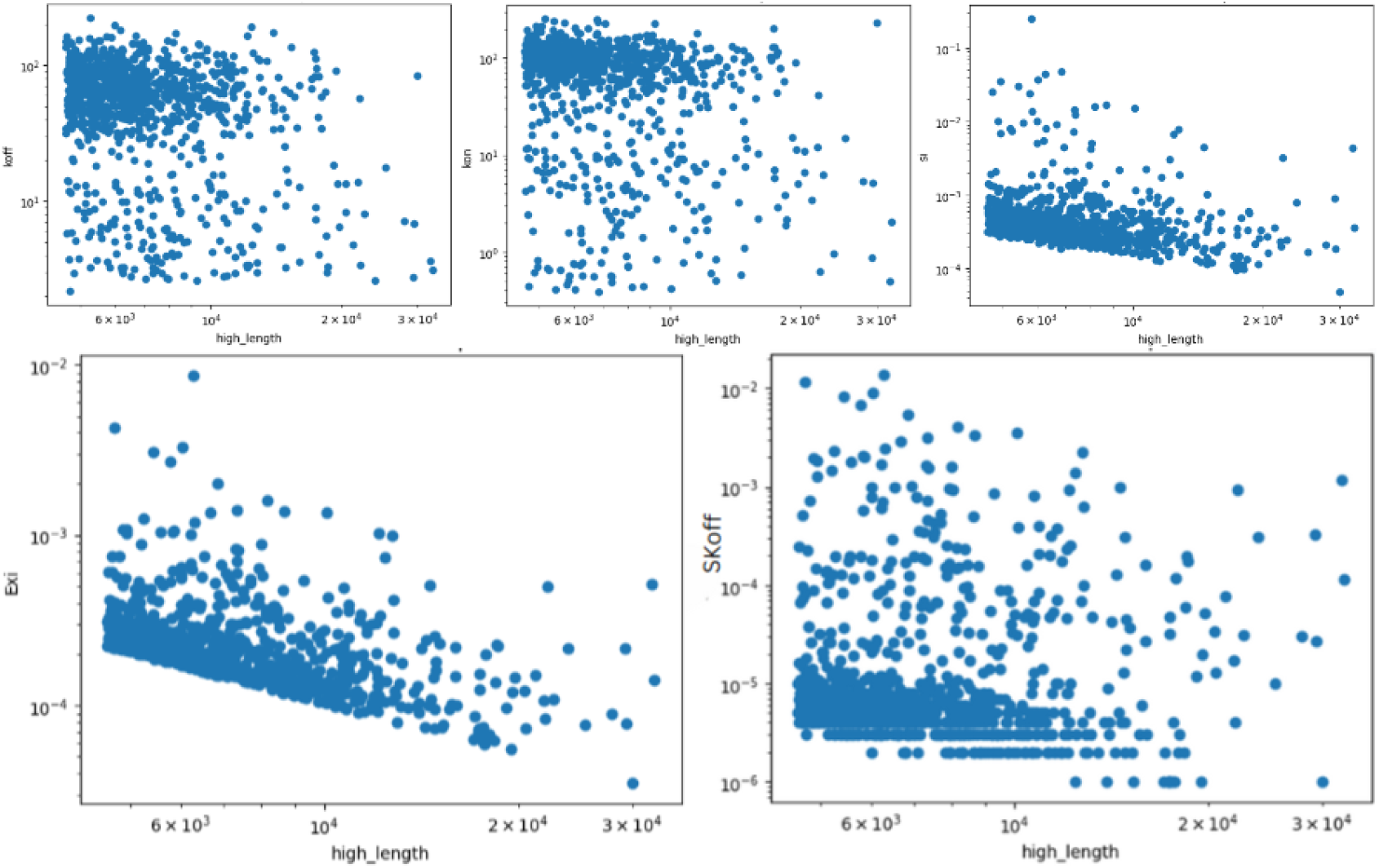
Relationship pattern between long length genes and kinetic parameters.

Since transcription is a time-dependent process, it makes sense that gene expression and gene length would be inversely correlated, and this is indeed the case ***Brown (2021)***.

We have seen from the figure 2, 3, *SK*_*off*_ has decreasing pattern for short length genes and it maks cluster for longer genes.

### Impact of various kinetic parameters on mean expression

Understanding the impact of various kinetic parameters on mean expression is essential for uncovering the mechanisms of gene regulation and transcription dynamics ***Raj and Van Oudenaarden (2008)***. Kinetic parameters such as *S*_*i*_, *K*_*on*_, *K*_*off*_, *SK*_*off*_ play critical roles in determining how genes are expressed under different conditions. By studying their influence, researchers can gain insights into the regulatory elements and factors that control gene expression ***Bintu et al. (2005)***. This knowledge is vital for improving predictive models of gene expression, which have applications in synthetic biology, pharmacogenomics, and disease modeling. Additionally, understanding these parameters helps elucidate the phenotypic consequences of changes in gene expression and how cells adapt to environmental changes. In therapeutic contexts, such as gene therapy and CRISPR gene editing, insights into kinetic parameters can inform strategies to achieve desired expression levels and target specific regulatory mechanisms. Overall, analyzing the impact of kinetic parameters on mean expression enhances our understanding of gene regulation and offers numerous applications in biology, medicine, and biotechnology.

### Selecting drug targets that require lower doses

Choosing drug targets that require lower doses of drugs is advantageous for several reasons. First, it reduces the likelihood and severity of side effects, which improves patient compliance and quality of life ***McCormack et al. (2011)***. Lower doses enhance the safety profile by minimizing toxicity risks, which is especially important for chronic treatments or in vulnerable populations. This approach also improves the therapeutic index, ensuring a larger margin between effective and toxic doses. Additionally, it decreases the risk of drug interactions, which is beneficial for patients on multiple medications. Economically, lower doses reduce manufacturing and distribution costs, making treatments more affordable and accessible. Furthermore, improved patient compliance with minimal side effects and simpler dosing regimens leads to better treatment outcomes. Finally, using lower doses can help minimize the development of resistance in the context of antibiotics and cancer therapies, contributing to more sustainable and effective healthcare practices. In our previous work, We have selected few proteins as target from single-cell transcriptomics data in asexual stages ***Choudhuri and Ghosh (2024)***. From these selected proteins, we have indentified one protein as drug target ***Choudhuri and Ghosh (2024)***. We have observed from our kinetic parameter results that our identified protein has higher burst size and higher mean value expression level compared to others selected proteins. Result has been given in table 2. We can see from the table 2, that *P F* 3*D*7_1246200 has higher burst size and higher mean value expression in all the stages. We have selected this protein as target protein in our previous research paper ***Choudhuri and Ghosh (2024)***. We can say from our result that we also need lower doses for *P F* 3*D*7_1246200 target protein.

**Table 1.** Error value of linear regression model.

| Stage | Error | Error excluding $S_i$ | Error excluding $K_{on}$ | Error excluding $K_{off}$ | Error excluding |
| --- | --- | --- | --- | --- | --- |
| Ring | 2.19 | 2.24 | 2.20 | 2.19 | 2.78 |
| trophozoite | 1.04 | 1.05 | 1.04 | 1.04 | 1.21 |
| Schizont | 4.30 | 5.74 | 4.40 | 4.31 | 2.63 |
| Gametocyte | 2.19 | 2.24 | 2.20 | 2.19 | 2.78 |

**Table 2.** Burst size and Mean value expression of selected proteins.

| protein name | Stage | Burst size $SK_{off}$ | Mean value expression $E_{xi}$ |
| --- | --- | --- | --- |
| *PF3D7_1038400* | Ring | 0.000001 | 0.000036 |
| *PF3D7_1038400* | trophozoite | 0.000001 | 0.000039 |
| *PF3D7_1038400* | Schizont | 0.000001 | 0.000038 |
| *PF3D7_1038400* | Gametocyte | 0.000001 | 0.000036 |
| *PF3D7_0810300* | Ring | 0.000625 | 0.000729 |
| *PF3D7_0810300* | trophozoite | 0.00009 | 0.000592 |
| *PF3D7_0810300* | Schizont | 0.001678 | 0.001874 |
| *PF3D7_0810300* | Gametocyte | 0.000625 | 0.000729 |
| *PF3D7_1105000* | Ring | 0.000092 | 0.000569 |
| *PF3D7_1105000* | trophozoite | 0.003531 | 0.003451 |
| *PF3D7_1105000* | Schizont | 0.007561 | 0.008474 |
| *PF3D7_1105000* | Gametocyte | 0.000092 | 0.000569 |
| *PF3D7_1246200* | Ring | 0.009469 | 0.003771 |
| *PF3D7_1246200* | trophozoite | 0.000333 | 0.001351 |
| *PF3D7_1246200* | Schizont | 0.005703 | 0.005834 |
| *PF3D7_1246200* | Gametocyte | 0.009469 | 0.003771 |

## Discussion

This study investigates the relationship between gene length and kinetic parameters of gene expression, focusing on their influence on mean expression levels. Our findings highlight the critical role of transcriptional kinetics, particularly the importance of burst size (*SK*_*off*_), *k*_*on*_ (activation rate), and *k*_*off*_ (inactivation rate) in modulating gene expression dynamics.

The results demonstrate that *SK*_*off*_, which denotes the burst size, significantly influences mean expression levels. The regression analysis indicated that excluding *SK*_*off*_ from the model substantially increased error rates, underscoring its pivotal role in determining gene expression. These findings align with previous studies emphasizing the regulatory capacity of transcriptional bursting in fine-tuning gene expression.

We observed distinct patterns in kinetic parameters between short and long-length genes. Short-length genes exhibited different transcriptional behaviors compared to long-length genes, suggesting that gene length influences kinetic regulation mechanisms. This observation provides a deeper understanding of how structural gene attributes can impact regulatory processes, potentially informing the design of synthetic biology applications.

The insights gained from this study have significant implications for drug discovery, particularly in selecting target genes for low-dose drug strategies. Genes with high burst size and expression levels are prime candidates for low-dose interventions due to their efficient transcriptional output and stable baseline activity. This approach not only enhances therapeutic efficacy but also minimizes side effects and resistance development, crucial in chronic and antibiotic treatments.

Overall, this research advances our understanding of gene expression regulation by elucidating the role of kinetic parameters in transcription dynamics. Future work could explore the application of these findings in therapeutic contexts, such as optimizing gene therapy strategies or developing CRISPR-based gene editing techniques. Further investigations using diverse cell types and conditions will help generalize these insights and pave the way for innovative applications in biotechnology and medicine.

## Methods and Materials

The standard two state stochastic kinetic model for gene expression posits that a gene toggles randomly between ‘on’ and ‘off’ promoter states, allowing mRNA transcription only in the ‘on’ state ***Larson (2011); Munsky et al. (2012)***. If a single rate-limiting step governs transcription rates and transitions between promoter states, the fluctuations can be modeled by a two-state Markov process with *k*_*on*_ denoting the activation rate and *k*_*off*_ the inactivation rate ***Larson (2011); Peccoud and Ycart (1995)***. The average waiting times in active and inactive states are described by 1/*K*_*off*_ and 1/*K*_*on*_. The average time for gene in a active state is 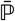 = 1/*K*_*off*_ /(1/*K*_*off*_ + 1/*K*_*on*_) = *K*_*on*_/(*K*_*on*_ + *K*_*off*_) ***Kim and Marioni (2013)***. Furthermore, it is assumed that the number of mRNA molecules in the gene decays at a rate of d per unit time and that transcription occurs at a rate of s per unit time while the gene is in the active promoter state ***Kim and Marioni (2013)***. Transcriptional bursting is defined by two parameters: burst frequency and burst size. Brust frequency is the frequency at which bursts occur per unit time (burst frequency, *k*_*on*_) and burst size or transcriptional efficiency (*s*/*K*_*off*_) is the average number of synthesized mRNA molecules during a gene’s active state. These four kinetic factors allow for deriving a set of differential equations that describe the time-dependent variation in the number of mRNA molecules (x) of a given gene within a cell ***Peccoud and Ycart (1995); Raj et al. (2006); Shahrezaei and Swain (2008)***. The steady-state solution of these equations has the following form:

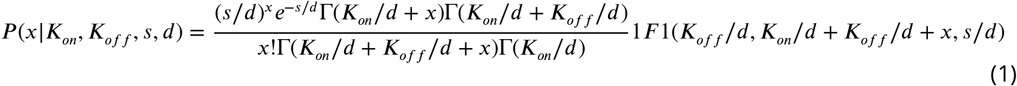

We have normalized equation 1 with d. 1/d is the average lifetime of an mRNA molecule. As we normalize all kinetic parameters with d, these will be time-independent. To simplify, we take the mRNA decay rate d equal to 1. However, computation of the confluent hypergeometric function is difficult because no computation method can reliably compute the function with all the parameters ***Kim and Marioni (2013)***. An auxiliary variable approach has been introduced to overcome this limitation ***Kim and Marioni (2013)***.

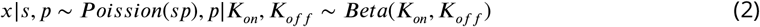

where p is a beta distribution-following auxiliary variable. The Poisson-beta distribution (PoBe), often known as the marginal distribution *P* (*x s, k*_*on*_, *k*_*off*_), has the same structure as the steady-state distribution given by equation 1 ***Kim and Marioni (2013)***. Given count measurements from RNAsequencing data, we assume that the number of reads mapped to a gene is proportional to the expression of the relevant mRNA molecule in the studied cell. As a result, the parameters of the kinetic model can be inferred using a Bayesian hierarchical approach, like a Gibbs sampler.

An auxiliary variable approach, introducing a Poisson-beta distribution, overcomes limitations and facilitates parameter estimation from RNA-sequencing data using Bayesian hierarchical methods. Yet, interpreting parameters remains a challenge, especially in identifying parameter space regions where transitions between promoter states are rapid or infrequent, influencing the distribution of mRNA molecules. To ensure statistical identifiability, fitting multiple models and employing goodness-of-fit statistics are recommended. Alternatively, hierarchical Bayesian modeling offers a comprehensive approach to determine the best-fitting distribution for each gene.

We have written the python code for this mathematical model. So, As a input if we give singlecell transcriptomics data to this model, it will predict the different kinetic parameters values for each gene.

### relationship pattern between gene length and other kinteic parameters

We have used single-cell transcriptomics data of plasmosium falciparum of asexual stages as input in our model and as a ouput result we got value of kinetic parameters of each genes. Next, we find the relationship between length and different kinetic parameters. We divded the genes into two categories, long-length genes and short-length genes. We find different relationship pattern between kinetic parameters and long-length, short-length genes. We have given pictures of patterns berween different kinetic parameters and low length genes in figure 2. Similarly we have given the pictures of patterns berween different kinetic parameters and high length genes in figure…. We can see that for low and high length genes kinetic parameters are following different kinds of pattern.

### Impact of various kinetic parameters on mean expression

Understanding the impact of various kinetic parameters on mean expression is essential for uncovering the mechanisms of gene regulation and transcription dynamics. Kinetic parameters such as *S*_*i*_, *K*_*on*_, *K*_*off*_, *SK*_*off*_ play critical roles in determining how genes are expressed under different conditions. By studying their influence, researchers can gain insights into the regulatory elements and factors that control gene expression. This knowledge is vital for improving predictive models of gene expression, which have applications in synthetic biology, pharmacogenomics, and disease modeling. Additionally, understanding these parameters helps elucidate the phenotypic consequences of changes in gene expression and how cells adapt to environmental changes. In therapeutic contexts, such as gene therapy and CRISPR gene editing, insights into kinetic parameters can inform strategies to achieve desired expression levels and target specific regulatory mechanisms. Overall, analyzing the impact of kinetic parameters on mean expression enhances our understanding of gene regulation and offers numerous applications in biology, medicine, and biotechnology.

We got our result after running our model and the result sheet contains kinetic parameter values of each genes. Next, a linear regression model has been trained on our result data and got error from this model. Our result dataset has six columns, Gene name, *S*_*i*_, *K*_*on*_, *K*_*off*_, *SK*_*off*_, *E*_*xi*_ and Our target column is mean vaue expression (*E*_*xi*_). Next, we have dropped one one column (lets say Si) and trained the same model on this reduced dataset. We observed that error value is increasing. We have dropped one column at a time one by one and noted error value for each time. We have seen that when we dropped *SK*_*off*_ and error increases significantly. We have done the same process for each stage and for each stage we have observed that if we dropped *SK*_*off*_ and error increases significantly.

### Selecting drug targets that require lower dose drugs

When considering target genes for low-dose drug strategies, it is essential to evaluate the interplay between burst size and mean expression level, which together determine a gene’s responsiveness to activation. Genes with high burst size and high expression level are ideal candidates for lowdose drugs, as they combine efficient transcriptional output per activation event with stable, high baseline activity, ensuring that minimal additional activation is required to achieve therapeutic effects. In contrast, genes with low burst size and high expression level may still be suitable targets if maintaining current expression levels is adequate, as their high baseline activity suggests significant involvement in cellular processes, though they may be less responsive to additional upregulation. Low burst size and low expression level genes typically require higher doses for significant upregulation, making them less desirable targets for low-dose strategies unless specific therapeutic contexts justify their modulation. Lastly, genes with high burst size but low expression level present an opportunity to harness their potential for substantial mRNA production during activation events, making them responsive to drugs that increase activation frequency, particularly in scenarios where induction of expression is necessary. These conditions offer a framework to prioritize genes based on their transcriptional dynamics and expression profiles, tailoring therapeutic strategies to achieve desired outcomes with minimal drug use.

## Acknowledgments

Additional information can be given in the template, such as to not include funder information in the acknowledgments section.

